# Integrating Diffusion and Liquid AI Models for Predicting Peptide Affinity from mRNA Display Selections

**DOI:** 10.64898/2026.05.05.723033

**Authors:** Colin M. Leaf, Pearl Qi, Yash Pragnesh Gandhi, Farzad Jalali-Yazdi, Justin N. Ong, Terry T. Takahashi, Rajiv K. Kalia, Richard W. Roberts

**Affiliations:** Department of Chemistry, University of Southern California, Los Angeles, California, 90089, USA; Mork Family Department of Chemical Engineering and Materials Science, University of Southern California, Los Angeles, California, 90089, USA; Thomas Lord Department of Computer Science, University of Southern California, Los Angeles, California, 90089, USA; USC Norris Comprehensive Cancer Center, University of Southern California, Los Angeles, California, 90089, USA; Thomas Lord Department of Computer Science, Department of Physics and Astronomy, Los Angeles, California, 90089, USA; Alfred E. Mann Department of Biomedical Engineering, University of Southern California, Los Angeles, California, 90089, USA

**Keywords:** protein design, mRNA Display, High Throughput Sequencing Kinetics, Machine Learning, Denoising Diffusion Implicit Network, Liquid AI, Liquid Time Constant Network, Closed-form Continuous Neural Network

## Abstract

*In vitro* selection and directed evolution technologies such as mRNA display, explore large libraries (≥10^14^ variants) and generate thousands to millions of functional polypeptide ligands to a variety of targets. Denoising diffusion implicit machine learning models (DDIMs) trained using display-derived deep sequencing data can greatly expand these functional sequences beyond what is accessible experimentally. However, methods are needed to predict peptide properties such as binding free energies (ΔG°). Here, we applied machine learning methods to predict binding free energies of both experimental and DDIM-generated peptide ligands against a target of interest, the oncogenic protein Bcl-x_L_. To do this, we trained a Closed-form Continuous (CfC) neural network using a dataset of 15,700 peptide ligands where pairs of sequences and their corresponding binding free energies (ΔG°) were used as inputs. This type of model was chosen due to its ability to represent irregular series. The resulting CfC model accurately predicts the rank order, within error, and binding free energies (ΔG°) for both experimental and DDIM-generated peptides, identifying five DDIM-generated peptides with single-digit picomolar affinities. Combining trained DDIM and CfC models offers a unified route to expand the scope of experimental ligand discovery, predict the molecular properties of both experimental and generated ligands, and highlights the utility of large quantitative datasets for making accurate *in silico* predictions of high-affinity peptide candidates.

**Statement:** High-throughput sequencing analysis of mRNA display libraries enables generating novel peptide ligands and expands the scope of functional sequences beyond what is accessible experimentally. Closed-form Continuous neural networks trained using sequences and their corresponding free energies accurately predict the binding free energies of both experimental and machine learning-generated peptides, enabling a route to quantitatively predict peptide properties using directed evolution data.

## 1. Introduction

*In vitro* selection techniques, such as phage, ribosome, and mRNA display, allow the discovery of novel peptides and proteins for diagnostics and therapeutics (Fan et al., 2024). Among these, mRNA display allows the search of libraries exceeding 10^14^ unique variants (Kamalinia et al., 2021; Mandecki, 1998; Roberts & Szostak, 1997). Recent advances in experimental methods, such as high-throughput sequencing kinetics (HTSK), enable the measurement of binding affinities at the pool level by coupling mRNA display with next-generation sequencing (Jalali-Yazdi et al., 2016). HTSK generates datasets comprised of tens of thousands of peptide sequences paired with their corresponding binding affinities (*K*_D_) or free energies (ΔG°).

Despite these advances, only a tiny fraction of the theoretical sequence space is experimentally accessible for proteins containing more than ∼10 randomized residues. To increase our ability to search more of sequence space, we have previously demonstrated that denoising diffusion implicit models (DDIMs) trained on high-throughput sequencing data from mRNA display selections (Figure S1) can generate novel, functional peptide sequences not present in the original selection pool (Qi et al., 2025). These generated peptides include functional sequences that were lost during experimental selection due to biases in synthesis, amplification, or non-binding steps, as well as sequences which were statistically improbable to be present in the starting library. A trained DDIM can trivially generate more than 10^5^ of these sequences, substantially expanding the searchable sequence space. However, while each sequence is likely functional, neither DDIM nor high-throughput sequencing can predict corresponding binding affinities for each generated sequence, making it difficult to choose which sequences to prioritize for experimental characterization.

Existing computational approaches have been used to predict peptide-protein interactions and binding affinities. In particular, machine learning approaches such as transformer-based protein language models, and structure prediction frameworks have enabled affinity prediction (Jumper et al., 2021; Rives et al., 2021; Romero-Molina et al., 2022; Zhou et al., 2022). While these methods have demonstrated strong performance, they can be computationally demanding and often require structural information, which is unavailable in many cases or can be experimentally challenging to obtain. Sequence-based deep learning frameworks that combine convolutional and attention mechanisms have also been developed to predict protein-peptide affinities and have shown impressive success across diverse targets (Sun et al., 2024). Such models are typically trained on broader, heterogeneous datasets spanning many proteins and sequence families, which can limit their sensitivity to the fine affinity differences among the homologous, high-affinity sequences commonly observed in selection datasets. As a result, their performance is reduced for sequences generated by models trained on selection results. Therefore, the challenge of predicting affinities for the large number of candidate peptides generated from high-throughput sequencing data remains an outstanding challenge.

Here, we used a Liquid AI-based predictive framework (R. M. Hasani et al., 2018) to predict peptide-target binding affinities (ΔG°) for both experimental and DDIM generated sequences. To do this, we combined deep sequencing data with quantitative binding constant measurements generated by high throughput sequencing kinetic analysis (HTSK) (Jalali-Yazdi et al., 2016). We selected this family of models because liquid AI models excel in fitting irregular series. For peptide-protein interactions, binding is governed by many factors, including the identity of each amino acid, its location in the sequence, and the surrounding residues. The relationship between a sequence and its binding free energy is thus nonlinearly distributed across sequence space. Further, experimental affinity measurements are typically unevenly distributed among the observed sequences, a feature which has proved challenging for traditional recurrent architectures to fit (Zong et al., 2025). Here, we explored treating peptide binding data as an irregular series wherein the encoded peptide sequence serves as the input signal and the binding free energy (ΔG°) serves as the system’s response (Figure 1). This approach results in robust, quantitative predictions using straightforward molecular encoding and does not rely on input structural information, giving a route to merge experimental and generative information toward the goal of predicting peptide properties.

**Figure 1:**
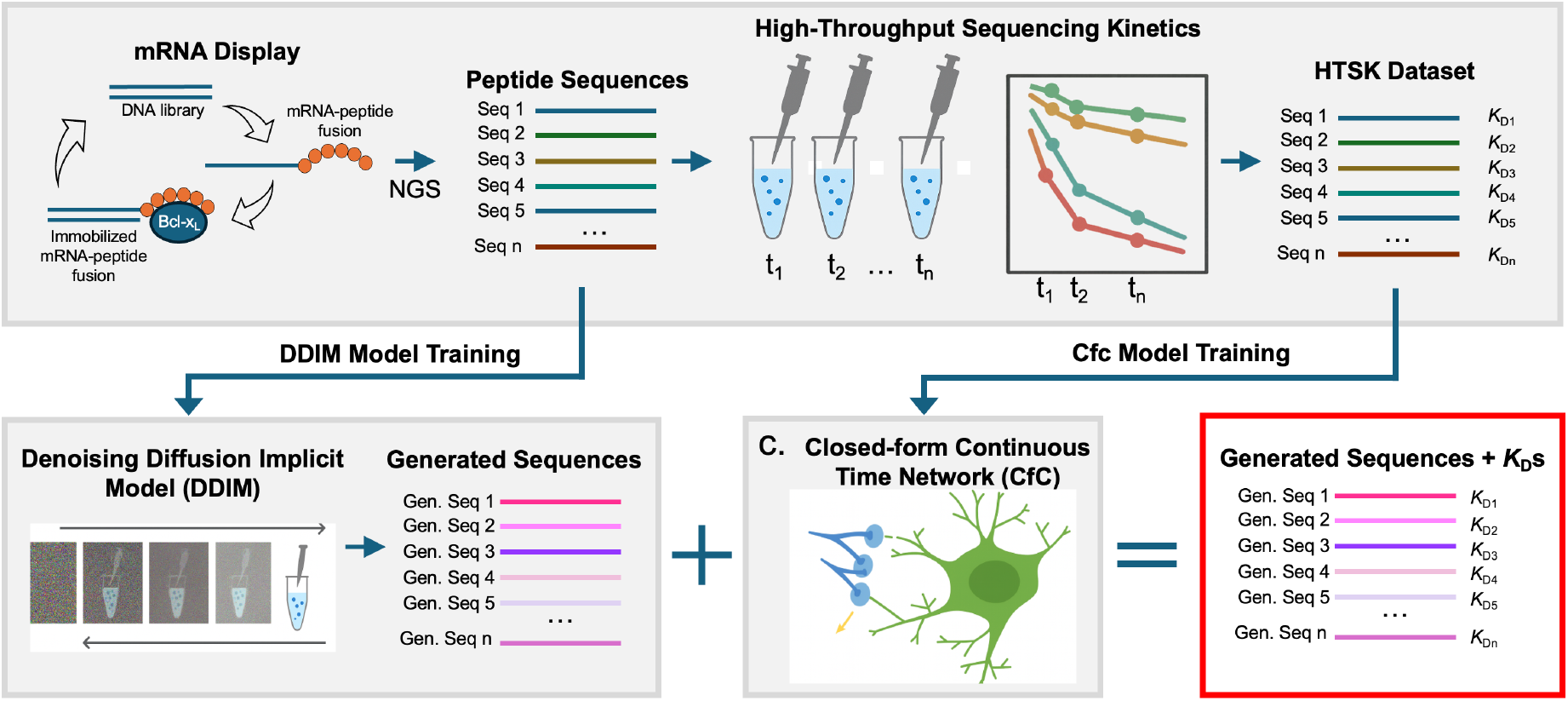
Pipeline for *In Silico* Generation and Evaluation of Novel Peptide Sequences. General schematic of the pipeline for the *in-silico* generation and characterization of novel proteins and peptides. First, high-affinity ligands are enriched from diverse libraries via mRNA display and off-rate selection. These ligands are identified and have their affinities characterized *en masse* via next generation sequencing and HTSK. The next generation sequencing data is used to train a DDIM, enabling the generation of novel sequences. The HTSK data is used to train a CfC, which learns to predict ligand affinity from the amino acid sequence. The trained CfC is then used to predict the affinity of the novel DDIM generated sequences, allowing for the identification of high affinity sequences not found in, or inaccessible through, the original library.

## 2. Results & Discussion

A long-term goal of our research is to generate novel, functional peptides by combining experimental and computational methods to predict and optimize peptide and protein properties. Previously, we had shown that deep sequencing datasets resulting from mRNA display directed evolution experiments could be analyzed via DDIM models. Once trained, these models can generate novel functional sequences far outside the scope accessible by mutation alone. However, these models do not provide quantitative metrics of function. Here, we worked to develop a unified framework for *in silico* peptide property prediction by combining HTSK, DDIMs, and liquid AI (CfCs) (Figure 1).

Liquid AI models belong to the family of the time-continuous neural networks that generalize recurrent architectures to differential systems (R. M. Hasani et al., 2018). The liquid time constant (LTC) network (R. M. Hasani et al., 2018) has demonstrated superior performance on irregular time series by incorporating adaptive time constants directly tied to the hidden states or the network, which allow the model to flexibly adjust to variations in input sequences. However, LTCs require numerical solvers during training, increasing training time and limiting their scalability. To overcome this, we employed the Closed-form Continuous (CfC) neural network (R. Hasani et al., 2022), which expresses the same dynamics in a differentiable closed form. The CfC introduces a sigmoid-based time-gating mechanism that efficiently balances memory and external input, achieving a 10-100X increase in training speed without compromising predictive accuracy.

Continuous-time models, such as CfCs, are adept at fitting irregular time series. We adapted this approach to use the peptide sequence and corresponding free energies as inputs. The training process effectively produces time constants that determine the response strength for any given amino acid change. For example, if a mutation of a particular amino acid results in a large change in affinity, the time constant in the model can be updated such that similar changes result in a strong response and vice versa for mutations with little impact on affinity. As the sequences are passed into the CfC and between its neurons, these neurons update their hidden states and time constants to minimize the difference between the model’s output and ground truth experimental values. This allows the model to learn a mapping which can predict the affinity of a sequence it has never encountered (Figure S2).

### 2.1 CfC Training and Evaluation Using High-Throughput Sequencing Kinetics Dataset

We began by training a CfC model using a previous dataset of 15,700 21-mer peptide sequences and their corresponding binding free energies. This dataset originated from an mRNA display selection of 21-mer peptides directed against the oncogenic protein B-cell lymphoma extra-large (Bcl-x_L_) (Jalali-Yazdi et al., 2014, 2015, 2016). Selection with an extension library yielded the high affinity peptide E1, and a subsequent doped selection yielded even higher affinity peptides. Of these 15,700 sequences, 10,990 were used for training, using mean squared error (MSE) as the loss function. During training, we observed a rapid decrease in MSE for both training and validation, indicating that the model was able to quickly learn a correlation between sequence and affinity and generalize that prediction across the diverse peptide set used for training (Figure 2a).

**Figure 2:**
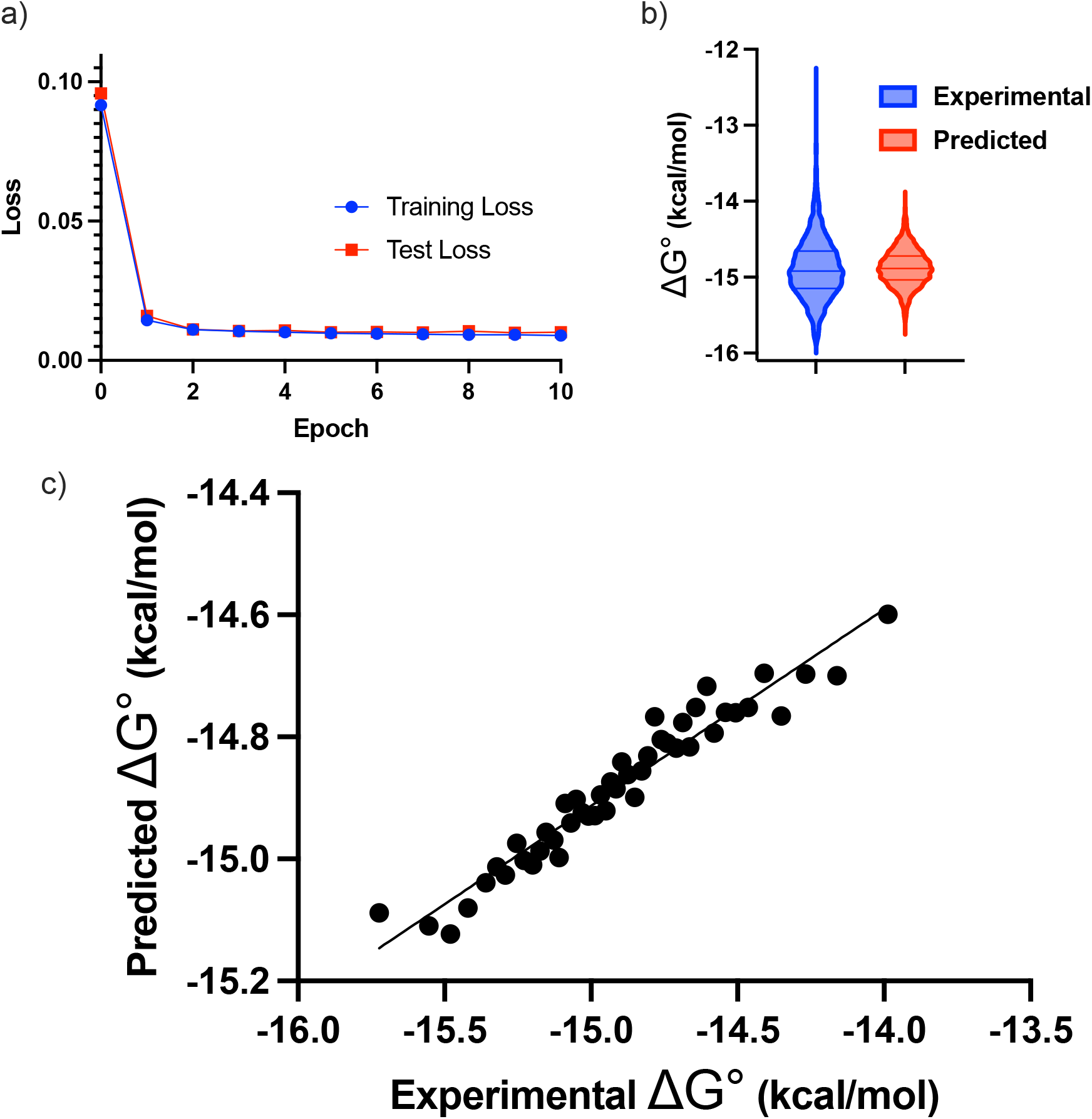
Evaluating CfC Performance on HTSK Dataset. (a) Training and validation loss curves for CfC model over 10 epochs. Both training and validation loss decrease rapidly within the first few epochs and converge to near-zero values, indicating fast learning and stable generalization. Mean Squared Error (MSE) is used as the loss function. **(b)** Violin plot comparing the distribution of experimental and predicted ΔG° (free energy) values. The predicted values closely follow the central tendency and spread of the true values, indicating that the model effectively captures the underlying distribution with only minor deviations. **(c)** Plot comparing the experimental and CFC predicted ΔG° values. Roughly, 2,300 test set sequences were grouped into sets of 50, and the groups’ median values were plotted. A strong correlation, and consistent bias are observed, demonstrating the model’s ability to capture the correlation between sequence and affinity.

We assessed the performance of the CfC model by comparing the predicted peptide binding affinity (ΔG°) values for the test set sequences to the ground truth experimental values obtained in the Bcl-x_L_ HTSK dataset (Figure 2b and 2c, Table 1). We observe that the distribution of predicted values closely mirrors that of the ground truth ΔG° values (Figure 2b). The predicted mean and median match both experimental values to the tenth of a kcal/mol (both -14.9 kcal/mol) (Table 1). The standard deviation of the predicted values (+0.22 kcal/mol) is only slightly lower than the true data (+0.49 kcal/mol), suggesting that the model exhibits slight smoothing but does not significantly underestimate variability (Table 1). The model also slightly compresses the free energy range of the dataset because nearly all sequences in the training set lie in the range of the model. However, this compression is not problematic as nearly all of the lost range is in the lower affinity range (i.e., weaker binders) which is of less interest.

**Table 1:**
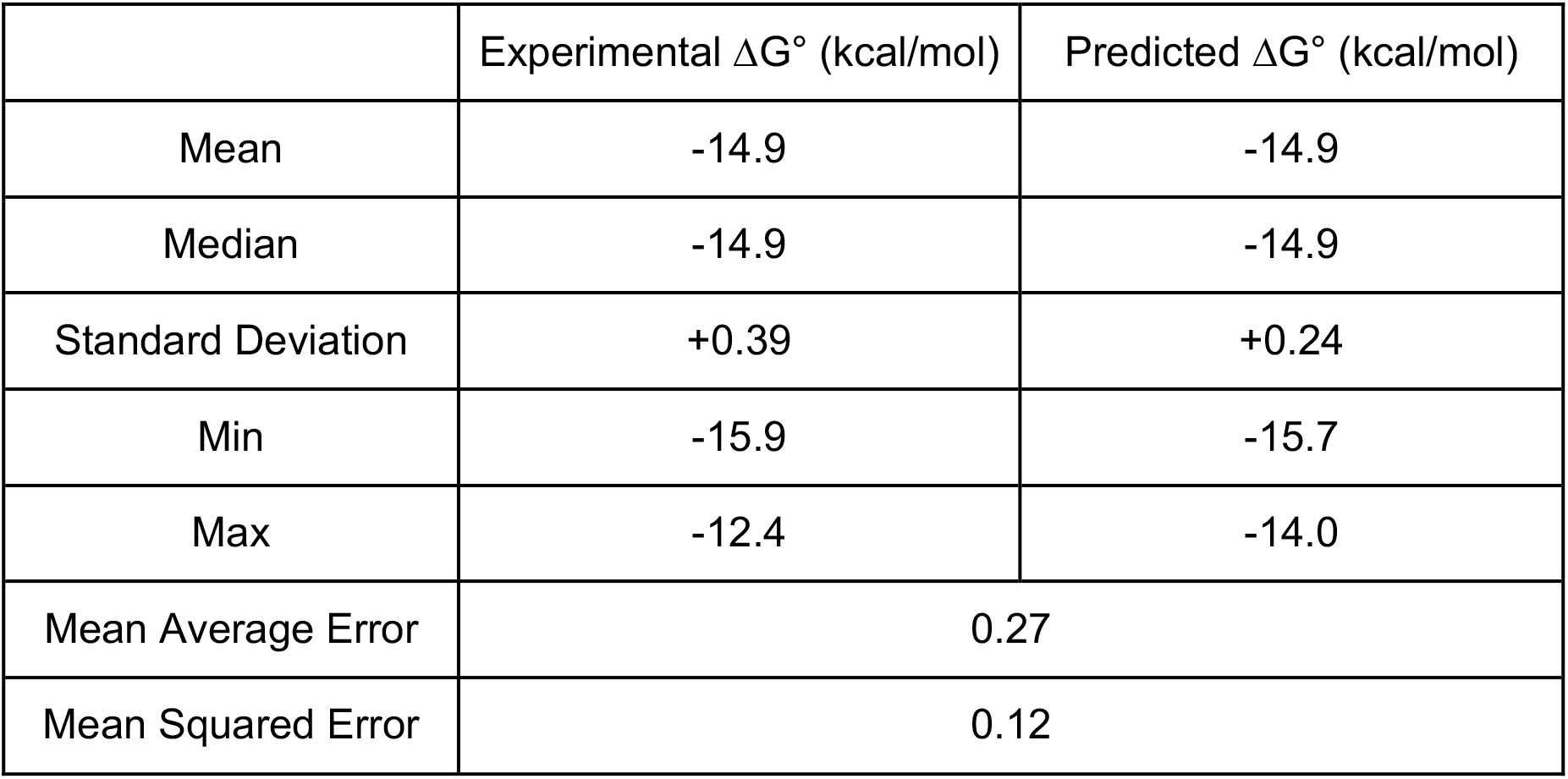
Key CfC Performance Statistics. Summary statistics of experimental (ground truth) and predicted ΔG° (free energy) values. The predicted values closely match the mean and median of the true distribution, with a slightly reduced standard deviation. The model predictions have a narrower range which is expected as a result of the dearth of experimental data in the lower affinity range.

| | Experimental $\Delta G^\circ$ (kcal/mol) | Predicted $\Delta G^\circ$ (kcal/mol) |
| --- | --- | --- |
| Mean | -14.9 | -14.9 |
| Median | -14.9 | -14.9 |
| Standard Deviation | +0.39 | +0.24 |
| Min | -15.9 | -15.7 |
| Max | -12.4 | -14.0 |
| Mean Average Error | 0.27 |  |
| Mean Squared Error | 0.12 |  |

We observe a strong correlation between experimental and predicted values across the dataset, with a consistent bias (Figure 2c). Impressively, the mean absolute error (MAE) and MSE are both significantly lower than +0.5 kcal/mol (Table 1). Together, these results demonstrate that the CfC model effectively generalizes across the training distribution and makes accurate predictions of free energy for unseen sequences from the HTSK dataset.

### 2.2 Predicting affinities for DDIM-Generated Sequences with CfC

Having successfully demonstrated that we were able to train the CfC to make affinity predictions for sequences chosen from within the HTSK dataset, we then sought to evaluate its performance predicting the affinities of novel peptides generated by DDIM. We first generated 20,000 novel sequences which do not appear in the HTSK dataset but share the same probability distribution. The similarity between the generated and training distributions was assessed by Kullback-Leibler divergence, yielding a value of 0.0036, indicating near-identical positional frequency distributions (Figure S4) (Kullback, 1951; Han, 2002). From these 20,000 novel sequences we selected four candidates (DDIM-HTSK1-4) spanning a range of binding affinities (Table 2). These sequences were then characterized experimentally to validate the performance of the CfC model. Predicted free energy values (ΔG°) were converted to dissociation constants (*K*_D_s) to facilitate interpretation in units of binding affinity.

**Table 2:**
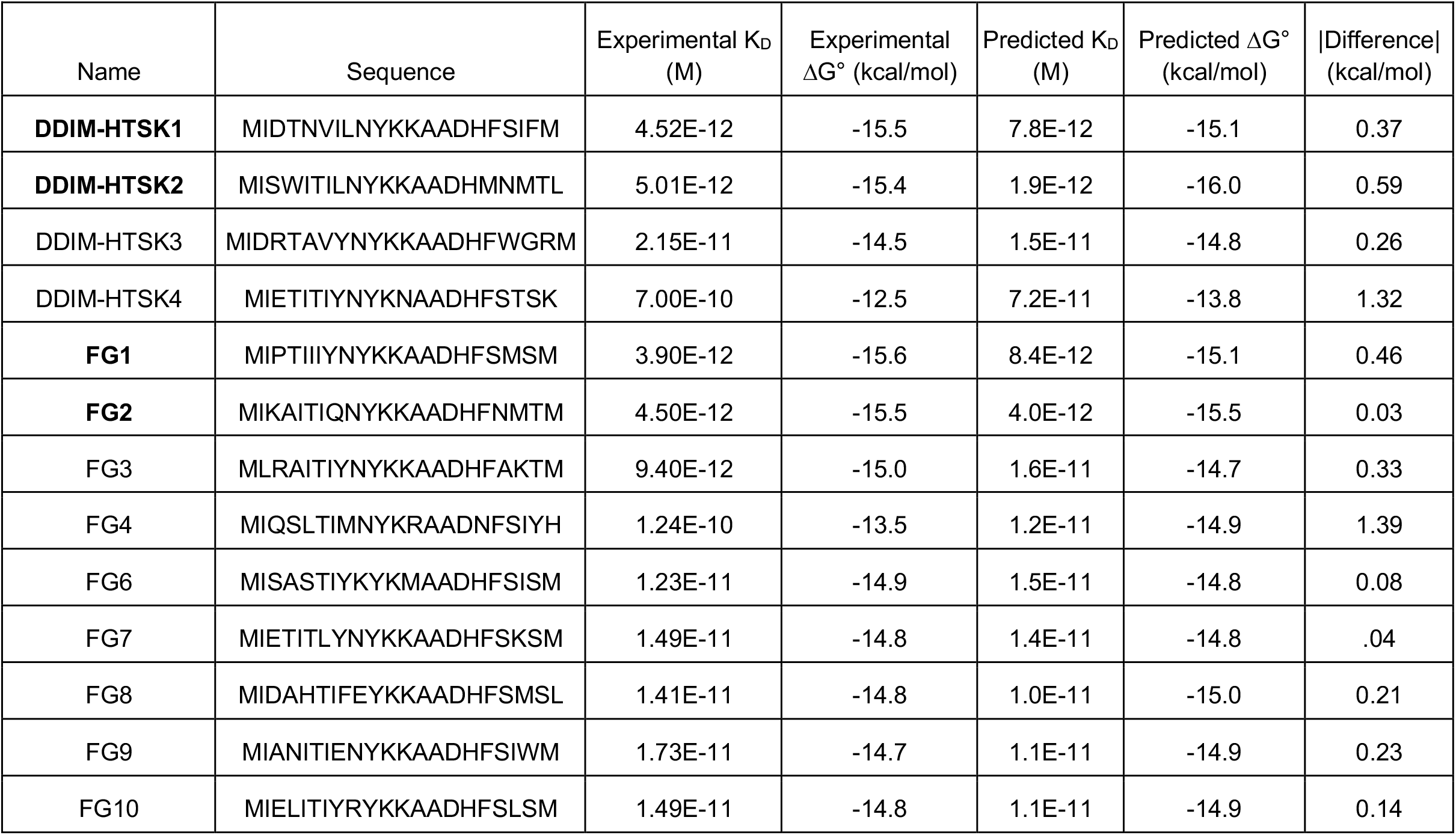
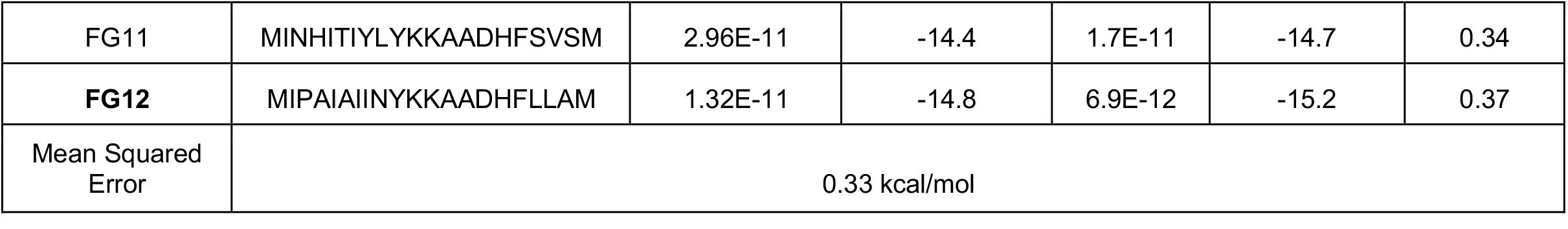
Experimental vs. Predicted Affinity for DDIM-Generated Sequences with High Predicted Affinities. Experimental vs. predicted affinity in ΔG° (kcal/mol) and *K*_D_ for the 14 kinetically evaluated DDIM generated peptides. For these sequences predictions were highly accurate with a MSE of less than 0.4 kcal/mol. Of the five sequences with the highest predicted affinity (bolded), four had a *K*_D_ ≤ 5 pM.

Among these candidates, the CfC model predicted DDIM-HTSK1 (MIETRTLFNYKKAADHFPRTK) and DDIM-HTSK2 (MIDPILIYNYKKAADDFSIGL) to have the strongest binding, with ΔG° values of -15.1 kcal/mol (8.4 pM) and -16.0 kcal/mol (1.8 pM), respectively (Table 2). HTSK3 (MIDRTAVYNYKKAADHFWGRM) was predicted to have a ΔG° value of -14.8 kcal/mol (15 pM), comparable to the wildtype sequence, E1 (-14.6 kcal/mol) on which the library founding the HTSK was based (Table 2). HTSK4 was predicted to have significantly weaker affinity (>70 pM) (Table 2).

Each sequence was synthesized as a radiolabeled mRNA-peptide fusion and after treatment with RNAse A the kinetic on- and off-rates of each sequence were measured at room temperature to match the temperature under which HTSK data was obtained. Experimental results confirmed the predictions made by the CfC. DDIM-HTSK1 and 2 had *K*_D_’s of 4.5 pM and 5.0 pM respectively, representing roughly four-fold improvements over the E1 peptide (Figure 3, Table 2). DDIM-HTSK3 (*K*_D_ = 21.5) showed similar affinity to the wildtype E1 (*K*_D_ = 18.4 pM) (Figure 3a-d, Table 2). DDIM-HTSK 4, which was predicted to exhibit substantially weaker affinity, demonstrated a relatively quick off-rate (Figure 3c) consistent with weaker overall affinity. Although all predicted *K*_D_ values fall within the picomolar range, it is important to note the predictive accuracy for lower-affinity sequences (such as DDIM-HTSK4) is expected to be significantly more variable. This is due to the composition of the training dataset, which is enriched for strong binders in the lower picomolar range and contains fewer examples of weaker binders in the higher picomolar range (Figure 3b). As a result, the model is optimized to distinguish and prioritize the more desirable high affinity binders (Lake & Baroni, 2017; Xu et al., 2020; Yu et al., 2024; Zhou et al., 2022). Despite this, our results highlight the ability of the CfC to accurately identify high affinity sequences generated by the DDIM, and to differentiate between high affinity and lower affinity candidates.

**Figure 3:**
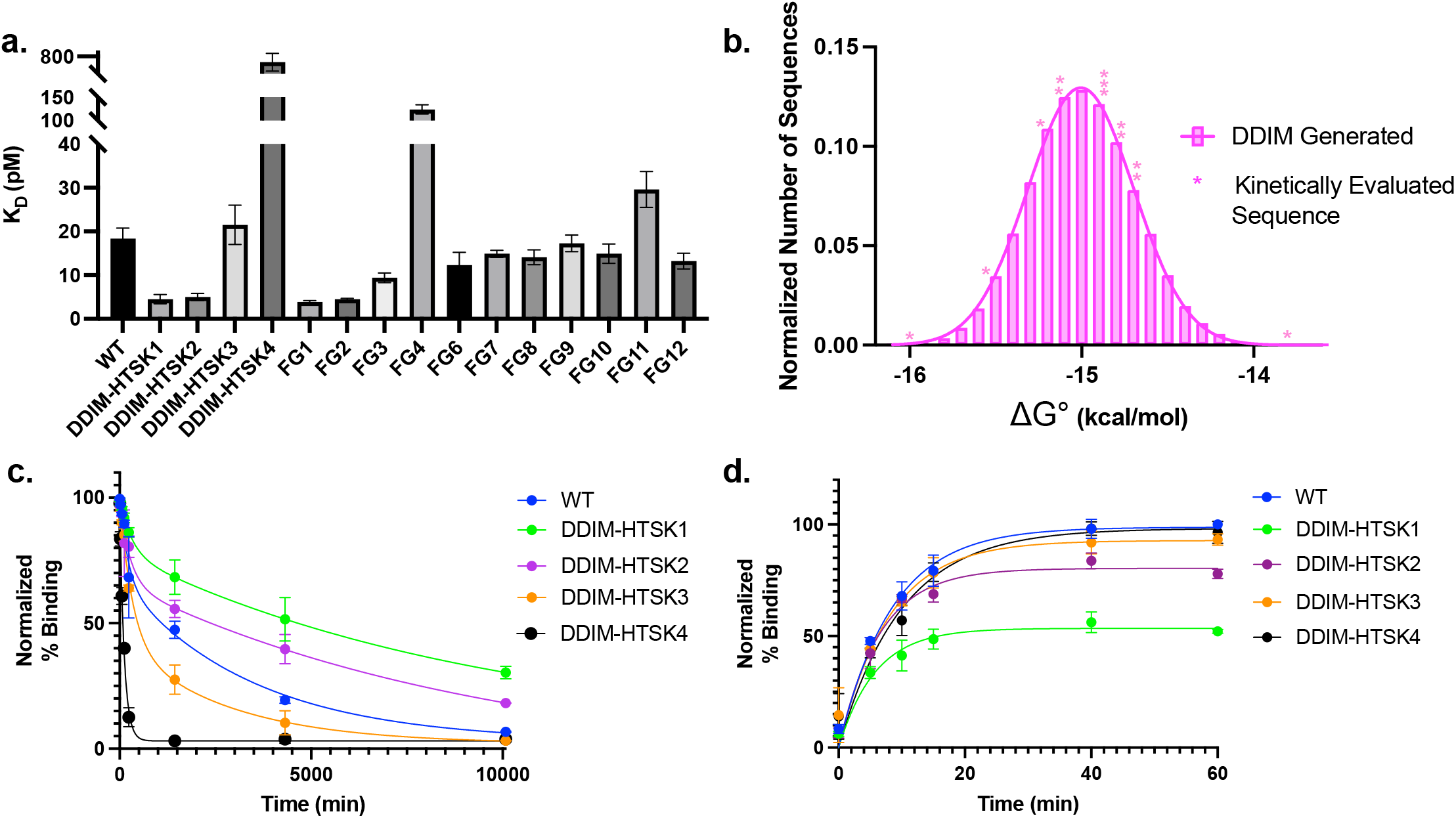
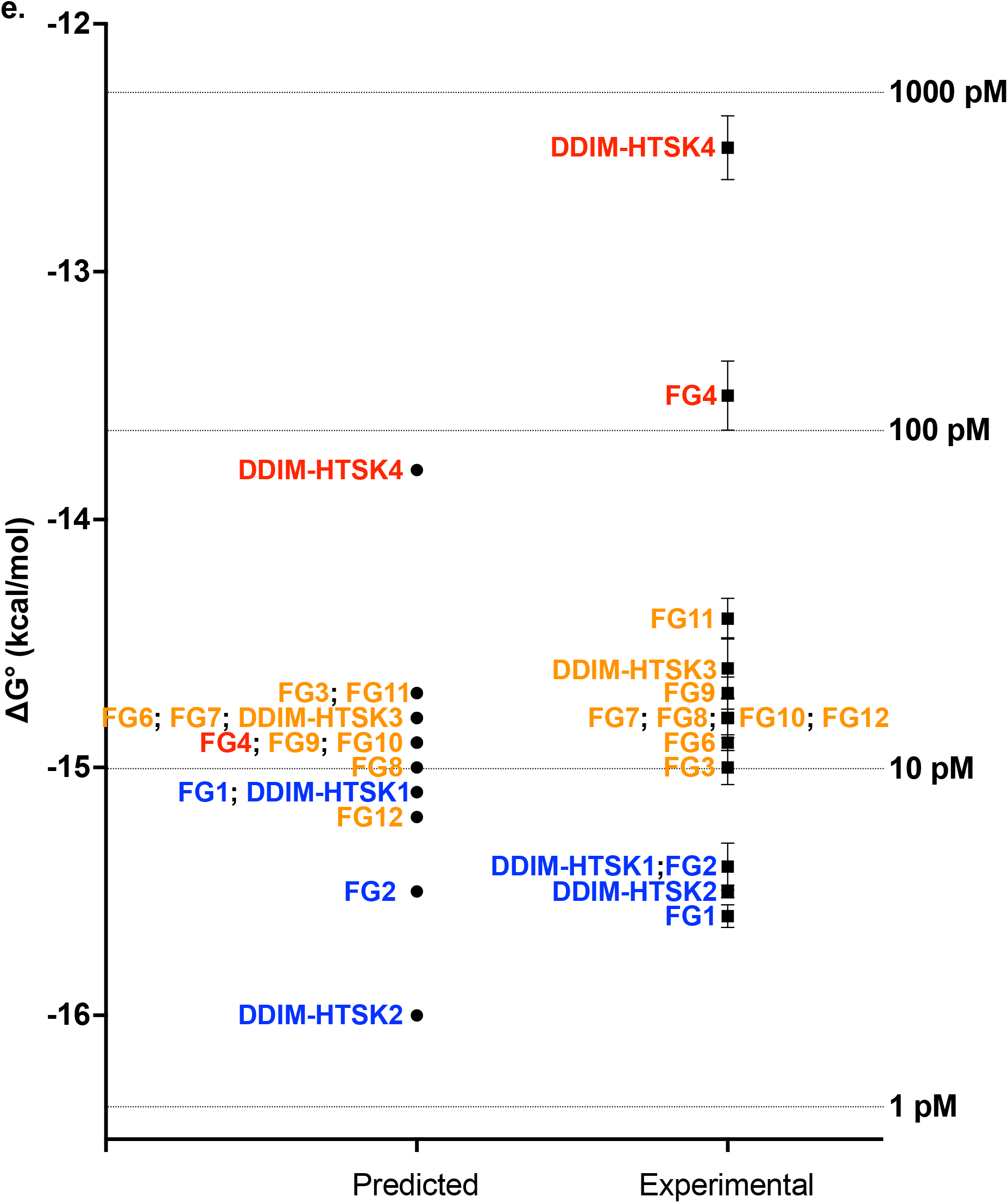
Kinetic Evaluation of DDIM Generated Peptide Predictions. **a)** Calculated binding kinetics (*K*_D_) of wildtype sequence E1 (WT) and DDIM generated peptides. Of the five generated peptides with the highest predicted affinities, four, DDIM-HTSK 1 and 2, and FG 1 and 2, have a *K*_D_ ≤ 5 pM **b)** Gaussian distribution of ΔG° predictions for DDIM generated sequences. Experimentally validated sequences are shown as stars at their predicted ΔG° **c)** Off-rate kinetics at room temperature for the four DDIM-generated peptides chosen as high affinity candidates, with the wildtype sequence E1 (WT) used as a control for comparison **d)** On-rate kinetics for the same four generated peptides, with binding at each time point normalized to the equilibrium binding of the wildtype sequences **e)** Predicted and experimental free energy values for DDIM generated peptides. Of the five sequences predicted to have the highest affinity, four had an experimentally determined *K*_D_ ≤ 10 pM. Additionally, DDIM HTSK4, predicted to be by far the lowest affinity binder, showed the highest experimental *K*_D_, demonstrating the model’s ability to identify and differentiate high and low affinity binders from DDIM generated sequences.

We next evaluated an additional 11 DDIM-generated sequences with high predicted affinities, FG1-4, and 6-12 (Table 2) (Qi et al., 2025). Analysis using the trained CfC predicted all ten of the sequences to be high-affinity binders (*K*_D_ < 20 pM) which matches well with the experimental data (Figure 3a). All sequences except two showed *K*_D_ values below 30 pM, and two of the three sequences with the highest predicted affinities had *K*_D_ values below 5 pM. Of particular note is FG7, which differs from DDIM-HTSK4 by only four amino acids. Despite this high degree of similarity, the model is able to correctly identify that FG7 is a high affinity binder but DDIM-HTSK4 is not, highlighting the ability of the model to accurately rank high and low affinity DDIM generated sequences.

The largest discrepancy between the CfC prediction and experimental results for the eleven predicted sequences was for FG4 (Figure 3e). This sequence is the most different of all tested peptides, with a hamming distance (Hamming, 1950) of nine from the nearest related experimental sequence in the training set. Overall, for these tested sequences the CfC was able to make accurate predictions, achieving a MSE of less than +0.4 kcal/mol (Table 2). Importantly, of the five tested sequences predicted to have the highest relative affinities, four had a *K*_D_ ≤ 5 pM, demonstrating that CfC based predictions are a suitable guideline for identifying high affinity DDIM generated sequences. While it is tempting to see this success and assume that sequences could be generated ad nauseum until increasingly high affinity binders are predicted and discovered, it is important to remember that the model is bounded by the dataset used to train it. Out of distribution detection (OOD) is an outstanding problem in machine learning, and consequently our model is able to identify high affinity peptides but is unable to separate peptides at the high affinity end of our distribution from peptides outside that distribution (Yu et al., 2024; Zhou et al., 2022). Thus the model makes the best predictions where there is high sampling and the goal of predicting DDIM generated binders with affinities far outside the measured training set, either high or low affinity, remains an open problem.

Reduced sequence sampling size can also limit predictions in low-complexity regions of the peptide ligands where sequence changes are sparse. Within the Bcl-x_L_ pool, there is almost no diversity between residues 10 and 15 (`Y-KAAD`) and within the original training dataset 99.9% of all sequences contain this exact sequence motif. We used DDIM to generate sequences that had changes in this region to test the limits of both the DDIM and CfC predictions. Four additional DDIM-generated sequences (DDIM-HTSK5-8) incorporated mutations that did not appear in the experimental selection data. For example, DDIM-HTSK5 contains a proline at position 18 that should disrupt the presumed alpha helical secondary structure of the binder (Woolfson & Williams, 1990). DDIM-HTSK6-8 contain one or more mutations within the conserved core motif (positions 13-15) (Table S1) that are also not seen in the experimental training set. While the trained CfC model predicts that each of these should have high affinity (*K*_D_ = 1.5 – 7 pM) all four sequences show little affinity in pull-down experiments (Supporting Information, Table S1). These erroneous DDIM predictions may result from two features: 1) the way a peptide sequence is encoded as an image matrix of amino acid vs. position, rather than with explicit molecular structure and 2) the very low sampling of alternative amino acids within this region. The error in the CfC predictions for these sequences is likely due to the small dynamic range of free energy differences used to train the model. The original pool contains only high affinity sequences such that the ΔG° values range over ∼ 2 kcal/mol. The CfC model may thus be unable to accurately represent predictions where point mutations produce large free energy changes without improved sampling of these sequences.

## 3. Conclusion

This work demonstrates that machine learning models can be trained using directed evolution data (pairs of sequences and free energies) to both generate new sequences and predict their binding free energies. The model requires a relatively large training set of binding free energies to produce accurate predictions, which we generated by high throughput sequencing kinetic analysis (Jalali-Yazdi et al., 2016). The CfC model achieved high accuracy on both experimental and DDIM-generated sequences (MSE < 0.4 kcal/mol for both), with several predicted peptides exhibiting four-to five-fold higher affinity than the wildtype. Importantly, these predictions do not require structural information as an input.

Combining the ability to generate new sequences using the DDIM with the ability to estimate their binding affinities using the CfC not only provides a powerful method for the discovery of new peptides, but also provides a route to optimize other physical properties of interest besides affinity. The DDIM can generate diverse sets of peptide ligands which will vary in physical properties of interest, such as hydrophobicity, net charge, and amino acid content. Because the CfC is able to estimate the binding affinities of such diverse peptides, these other important physical properties can be optimized concurrently with affinity. This provides a method to address some of the multiple physiochemical criteria that a ligand often needs, for example during optimization of a lead compound in therapeutic development.

In addition to binding constant prediction, the CfC model could be applied to make predictions of any quantifiable physiochemical property that is encoded by a peptide sequence, such as protease resistance or cell permeability. Future work will focus on peptide optimization of multiple properties simultaneously, accelerating functional peptide design across diverse targets of interest.

## 4. Material and Methods

### 4.1 High-Throughput Sequencing Kinetic (HTSK)

HTSK experiments were performed as previously described (Jalali-Yazdi et al., 2016). Briefly, radiolabeled mRNA-peptide fusions were reverse transcribed and incubated with Bcl-x_L_ immobilized on magnetic beads under standard binding conditions. Aliquots were collected at multiple time points during the association and dissociation phase, and bound fractions were isolated by magnetic separation. Radioactivity on beads was measured by scintillation counting and determining pool peptide remaining. Recovered DNA was PCR-amplified and subjected to Illumina sequencing. Sequencing counts of each peptide were multiplied by the total on-bead counts at each time point to obtain normalized abundance values. Normalized counts were used to calculate the on- and off-rate for individual peptides.

### 4.2 Training of the Diffusion Denoising Implicit Model and Generation of Novel Sequences

Each sequence was first converted into a 20 x 20 matrix of 0’s and 1’s through one-hot encoding (Rumelhart et al., 1986), with the starting methionine (M) removed prior to both training and generation. DDIM training was performed as previously described (Qi et al., 2025). Briefly, sequencing data from mRNA display selection against Bcl-x_L_ were used to train a Denoising Diffusional Implicit Model (DDIM). Peptide sequences were one-hot encoded as 20x20 binary matrices and input into the model. Noise was iteratively added over 50 timesteps, and the model was trained for 5,000 steps until reaching minimal mean square error. After training, the model was used to generate 20,000 novel peptide sequences, which were decoded to amino acid format for downstream analysis. The similarity between the generated and training distributions was assessed by Kullback-Leibler divergence, yielding a value of 0.0036, indicating near-identical positional frequency distributions.

### 4.3 Training of Closed-form Continuous (CfC) Model

A dataset of 15,700 peptide sequences and their corresponding binding free energies (ΔG°) obtained from high-throughput sequencing kinetics (HTSK) experiments was used for CfC training. ΔG° values were min-max normalized based on the training subset and rescaled after prediction. Peptides were encoded as 20x20 matrices and divided into training, validation, and test sets (70:15:15).

Each tensor was passed through a batch normalization layer, a 64-unit dense layer with *tanh* activation, a CfC recurrent block returning the full sequence, a 32-unit dense layer with *tanh* activation, and a second CfC block returning the final state. A linear dense layer produced the ΔG° prediction. The model was trained using the Adam optimizer with mean-squared error loss for 10 epochs. Training and validation losses were monitored over 10 epochs and converged within the first few epochs, indicating stable learning. Predicted ΔG° values were compared with experimental measurements in HTSK.

### 4.4 mRNA display for individual peptides of interest

ssDNA encoding peptide sequences of interest were ordered from Integrated DNA Technologies and PCR-amplified using Taq DNA polymerase. Each DNA construct contained a T7 promoter, a transcription start codon, an open reading frame, and a 3’ constant sequence (5’ – GGTTCAGGCAGTGGT – 3’). The PCR-amplified DNA was transcribed in a transcription buffer (80 mM HEPES-KOH, pH 7.5, 2 mM spermidine, 40 mM DTT, 25 mM MgCl_2_). The mixture was briefly heated before initiating by the addition of T7 RNA polymerase. Reactions were incubated at 37 °C overnight and quenched with 0.1 volume of 0.5 M EDTA, pH 8.0. The resulting mRNA was ligated to a puromycin-DNA linker (F30P) using a splint oligonucleotide (5’ – TTTTTTTTTTTTACCACTGCCTGA – 3’) and T4 DNA ligase in a ligation buffer (50 mM Tris-HCl pH 7.5, 10 mM MgCl_2_, 1 mM ATP, 10 mM DTT) for 60 minutes at room temperature. Following ligation, the product was gel-purified by urea-PAGE, recovered, and concentrated. The sample was then desalted and concentrated using a 0.5 mL 30k MWCO Amicon centrifugal filter. Translation was carried out in rabbit reticulocyte lysate with [^35^S]-methionine to yield mRNA-peptide fusions, which were purified with oligo-dT-agarose beads. After washing, the mRNA-peptide fusions were eluted in hot water.

### 4.5 4Equilibrium binding and binding constant determination

Radiolabeled mRNA-peptide fusions were incubated with Bcl-x_L_ immobilized on Neutravidin agarose beads in selection buffer (1x PBS, 0.025% (v/v) Tween-20, 0.1% (w/v) BSA) for 60 minutes at 4 °C. After incubation, beads were washed three times with a blocking buffer (selection buffer and 20 µM biotin) to remove unbound peptides. The supernatant, washes, and beads were counted in a scintillation counter.

For association kinetics, the displayed peptides were first mixed with Bcl-x_L_ immobilized on Neutravidin beads. Aliquots were removed at defined time points (1, 5, 10, 15, 40, 60 minutes). Beads and supernatants were separated and quantified by scintillation counting. For dissociation kinetics, the displayed peptides were incubated with the target beads for 60 minutes, washed, resuspended in blocking buffer (selection buffer and 20 µM Biotin), and challenged with a 100-fold excess of non-biotinylated Bcl-x_L_. Aliquots were collected at various time points to measure radioactivity and determine the percentage of remaining peptide. All experiments were performed in technical triplicate (n = 3). The on-rate was determined using an association kinetic model, with a single concentration of the radiolabeled peptide. The off-rate was determined using a dissociation one-phase decay model using GraphPad Prism 10.0.

## Acknowledgements

This work was performed, in part, at the Center for Integrated Nanotechnologies, an Office of Science User Facility operated for the U.S. Department of Energy Office (DOE) of Science. Sandia National Laboratories is a multimission laboratory managed and operated by National Technology & Engineering Solutions of Sandia, LLC, a wholly owned subsidiary of Honeywell International, Inc., for the U.S. DOE’s National Nuclear Security Administration under Contract DE-NA-0003525. This work was also supported by NIH grants R01CA170820 (R.W.R. and T.T.T.), R01NS125769 (R.W.R.), and R21GM144910 (T.T.T.). This work was also supported by The Los Angeles Rubber Group Inc. (TLARGI), which provided fellowship funding to P.Q. This work was supported by the Center for Advanced Research Computing (CARC) at the University of Southern California, which provided computing resources that contributed to the research results reported within this publication, the USC Molecular Genomics Core (supported by the Norris Comprehensive Cancer Center CCSG grant (NCI grant# P30CA014089)), the USC Agilent Center of Excellence in Biomolecular Characterization, and the USC Center for Peptide and Protein Engineering.

