## Supplementary material for "Integrating Diffusion and Liquid AI Models for Predicting Peptide Affinity from mRNA Display Selections": Figure S1, S2, S3, S4, and Table S1

Supplemental

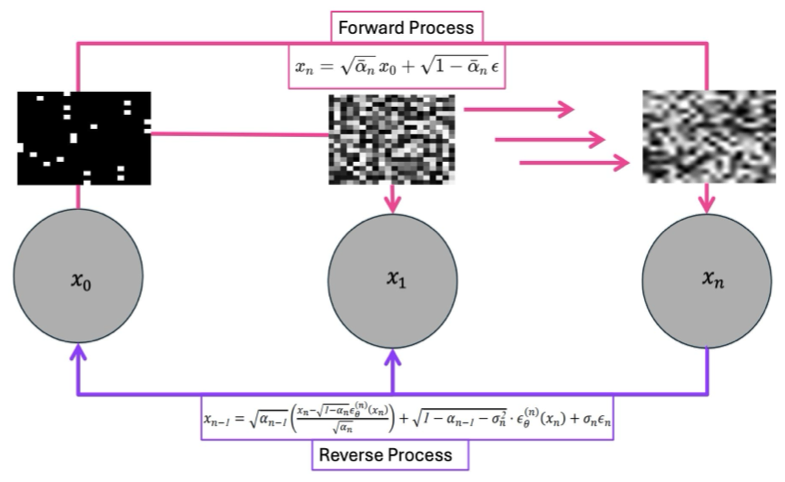

Figure S1. Diagram of a DDIM applied to the generation of novel peptide sequences (Qi et al., 2025; Song et al., 2020). Peptides are first converted into black and white images via one-hot encoding (Rumelhart et al., 1986). These images are then used to train the DDIM, which can then produce novel images sampled from the same probability distribution.

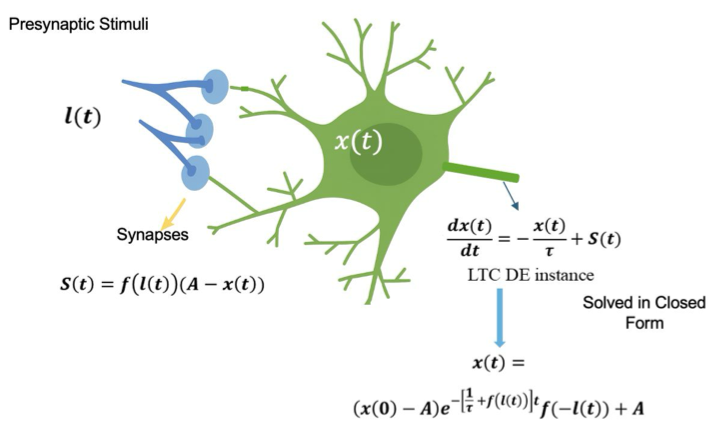

Figure S2. Closed-form approximation of nonlinear synaptic integration in an LTC unit.

A postsynaptic neuron receives input I(t) through a nonlinear conductance-based synapse, producing current S(t) that drives membrane potential x(t). The LTC differential equation governing x(t) is approximated here in closed form to illustrate the interaction between synaptic input and neuronal dynamics.

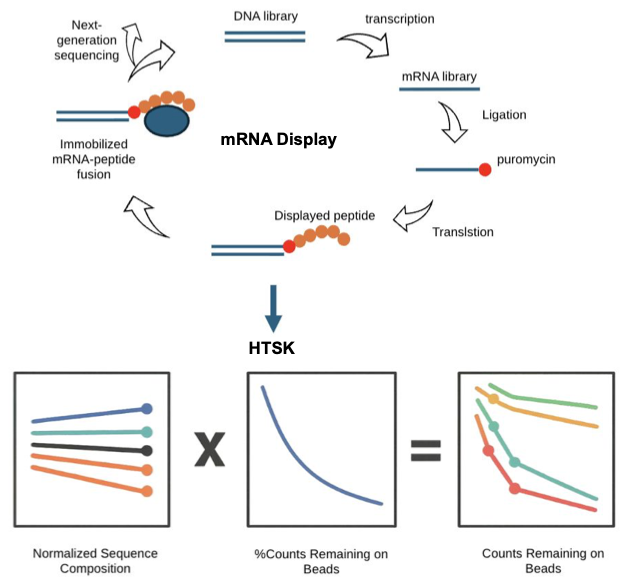

Figure S3. Diagram of mRNA display and high-throughput sequencing kinetics (Jalali-Yazdi et al., 2016). First, several rounds of selection are performed, enriching the pool in high affinity sequences. Once a pool of the desired affinity is obtained, HTSK is performed to identify peptide sequences and corresponding affinities *en masse*.

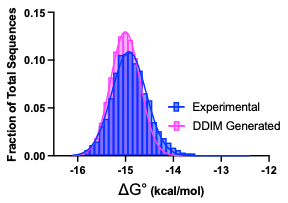

Figure S4. Overlay of sequence distribution for DDIM generated sequences and test-set experimental sequences. Differences between the two distributions are small, as demonstrated by a low Kullback-Leibler divergence of 0.0036.

| Name | Sequence | Experimental ∆G° (kcal/mol) | Predicted *K*_D_ (M) | Predicted ∆G° (kcal/mol) |
| --- | --- | --- | --- | --- |
| DDIM-HTSK5 | MIETIWIYLYKKAADHF**P**AGM | NA | 6.97E-12 | -15.2113285 |
| DDIM-HTSK6 | MIDWIIIKNYKKAA**R**HFNMFI | NA | 1.46E-12 | -16.138863 |
| DDIM-HTSK7 | MIDPITIQNYKMAA**F**HFYMSI | NA | 2.64E-12 | -15.786223 |
| DDIM-HTSK8 | MISQVSILQYKR**C**A**H**HFFMSL | NA | 4.00E-12 | -15.540102 |

Table S1. DDIM generated sequences (DDIM-HTSK5-8), which were predicted by the CfC to be high affinity, but contain changes in the conserved core motif. Mutations which were expected to be severely deleterious and do not appear in the training set are in bold.
